# Staurosporine drives non-canonical melanocyte maturation by coupling β-catenin signaling to actin-dependent dendrite remodeling

**DOI:** 10.64898/2026.06.05.730514

**Authors:** Kangning Xu, Lingli Yang, Sylvia Lai, Fei Yang, Yasutaka Kuroda, Daisuke Tsuruta, Ichiro Katayama

## Abstract

Skin pigmentation relies on the coordinated regulation of melanin production and dendritic morphology to ensure effective pigment distribution. While staurosporine is widely used as a proapoptotic agent in malignant cells, its effects on normal human melanocytes have not been fully characterized. Here, we investigated the impact of staurosporine on melanocyte biology and identify it as a potent inducer of non-canonical melanocyte maturation at sub-cytotoxic concentrations. In primary human neonatal melanocytes, staurosporine treatment enhances melanogenesis and promotes pronounced dendritic remodeling, leading to functional maturation distinct from its apoptotic effects in melanoma cells. Phenotypic analyses demonstrate increased pigment production and expanded dendritic networks that support efficient pigmentation. Molecular characterization indicates that these effects are associated with coordinated activation of β-catenin signaling and actin-dependent cytoskeletal remodeling. The physiological relevance of these findings was further examined in vivo. Topical application of staurosporine to normal guinea pig skin increased baseline pigmentation without detectable inflammation. In addition, staurosporine accelerated repigmentation in a rhododendrol-induced leukoderma model by restoring functionally mature melanocyte populations and enhancing nuclear localization of β-catenin. Together, these results identify staurosporine as a non-canonical modulator of melanocyte maturation and highlight the coordinated regulation of pigment production and dendritic remodeling as a key process supporting pigmentation in acquired hypopigmentary conditions.

**Summary:**

- Staurosporine promotes non-canonical maturation of human melanocytes at sub-cytotoxic concentrations.
- Treatment enhances both melanogenesis and dendritic remodeling, supporting functional pigmentation.
- Staurosporine increases baseline skin pigmentation *in vivo* without inducing inflammation.
- Repigmentation is accelerated in a rhododendrol-induced leukoderma model through restoration of mature melanocyte populations.
- These findings highlight coordinated regulation of pigment production and dendritic morphology as a potential strategy to promote pigmentation in acquired hypopigmentary conditions.

**Significance:** Loss of melanocyte dendricity and functional maturation is a shared feature of multiple acquired hypopigmentary disorders, including vitiligo and chemical-induced leukoderma. This study demonstrates that staurosporine promotes dendritic remodeling and pigmentation in normal human melanocytes and enhances repigmentation in vivo. By identifying a melanocyte-intrinsic, ultraviolet-independent maturation program, our findings provide a biological framework for strategies aimed at restoring functional melanocytes in depigmented skin.

## 1. Introduction

Melanocytes are specialized neural crest–derived cells located in the basal layer of the epidermis, where they synthesize melanin within melanosomes and transfer these organelles to surrounding keratinocytes. This process relies on an elaborate dendritic network that enables efficient melanosome transport and distribution, thereby determining skin pigmentation and contributing to photoprotection against ultraviolet radiation (Lin & Fisher, 2007; Westerhof, 2006). Accordingly, melanocyte dendricity is not merely a morphological characteristic but a critical functional determinant of the epidermal melanin unit (Fitzpatrick & Breathnach, 1963; Lin & Fisher, 2007). Disruption of dendritic architecture frequently precedes visible pigment loss and is a shared feature of multiple hypopigmentary conditions, including vitiligo, post-inflammatory hypopigmentation, and chemically induced leukoderma (Lin & Fisher, 2007). Despite its central importance, regulatory mechanisms that coordinate melanocyte dendritic remodeling with pigmentation remain incompletely understood.

Staurosporine is a microbial alkaloid originally characterized as a potent, broad-spectrum kinase inhibitor (Omura, Sasaki, Iwai, & Takeshima, 1995; Tamaoki et al., 1986) and has long been used as a canonical inducer of apoptosis in melanoma and other cancer cells (Bertrand, Solary, O’Connor, Kohn, & Pommier, 1994; Gescher, 1998; Zhang, Gillespie, & Hersey, 2004). At sub-cytotoxic concentrations, however, staurosporine has been reported to exert diverse biological effects in non-malignant cells, particularly in neuronal lineages, where it promotes neurite outgrowth (Kohno, Ninomiya, Kikuchi, Konno, & Kojima, 2015; Maeda, Tomita, Fukuda, & Tagami, 1992; Sano, Iwanaga, Fujisawa, Nagahama, & Yamazaki, 1994; Thompson & Levin, 2010), morphological differentiation (Borge, Lemare, Demignot, & Adolphe, 1997; Rasouly et al., 1996; Thompson & Levin, 2010), and cytoskeletal reorganization (Borge et al., 1997; Hoben & Athanasiou, 2008; Kim, Song, Jin, & Sonn, 2012; Kohno et al., 2015; Sano et al., 1994). These observations suggest that staurosporine can influence cell shape and differentiation programs in a context-dependent manner. Whether similar non-apoptotic, morphogenetic effects occur in normal human melanocytes has not been systematically examined.

Here, we report that staurosporine elicits fundamentally distinct responses in normal melanocytes compared with melanoma cells. While melanoma cells undergo apoptosis in response to staurosporine, primary human melanocytes exhibit marked resistance at comparable concentrations. Instead, staurosporine robustly enhances melanocyte dendricity, pigmentation, and functional maturation. These effects are associated with coordinated regulation of cytoskeletal dynamics and melanogenic activity, indicating activation of a non-canonical maturation program. We further demonstrate the physiological relevance of this response *in vivo*, as topical staurosporine application increases pigmentation in guinea pig skin and accelerates repigmentation in a rhododendrol-induced leukoderma model. Together, our findings identify staurosporine as a powerful modulator of melanocyte morphology and pigmentation and reveal an intrinsic pathway linking dendritic remodeling to pigment production.

## 2. Materials and Methods

### 2.1. Reagents

Staurosporine (purity > 99%) was obtained from Cell Signaling Technology, Inc. (CST; Product No. 9953; Danvers, MA, USA). For *in vitro* experiments, staurosporine was dissolved in DMSO. For *in vivo* experiments, it was formulated in a 50% ethanol/50% sesame oil (*v/v*) mixture. For all *in vitro* treatments, DMSO concentrations were adjusted to a final concentration of 0.1% (*v/v*) in the growth medium. RD (4-(4-hydroxyphenyl)-2-butanol; trade name Rhododenol®) was generously provided by Kanebo Cosmetics Inc. (Tokyo, Japan). A 20% (*w/v*) rhododendrol stock solution was prepared in a 50% ethanol/50% sesame oil (*v/v*) mixture.

### 2.2 Cell lines and cell culture

HEMn-MPs (normal human primary epidermal melanocytes derived from moderately pigmented neonatal foreskin) were obtained from Invitrogen (Carlsbad, CA, USA) and cultured in medium 254 (M-254-500; Thermo Fisher Scientific) supplemented with 1% (*v/v*) human melanocyte growth supplement. G-361 human melanoma cells were obtained from the Japanese Collection of Research Bioresources (Osaka, Japan) and maintained in high-glucose Dulbecco’s modified Eagle medium supplemented with 10% fetal bovine serum and 1% penicillin-streptomycin (Thermo Fisher Scientific, Waltham, MA, USA). All cultures were maintained at 37°C in a humidified incubator with 5% CO_2_.

### 2.3. RNA interference

For siRNA-mediated knockdown of β-catenin, GSK3β, CDC42, and Rac1, HEMn-MPs were transfected with 30 nM predesigned Silencer® Select siRNAs (si-β-catenin: s436; si-GSK3β: s6240; si-CDC42: VHS40396; si-Rac1: s11712; Thermo Fisher Scientific) using Lipofectamine® RNAiMAX (Invitrogen), in accordance with the manufacturer’s instructions.

### 2.4. Cell viability assay

HEMn-MPs and G-361 cells (2.5 × 10^4^ cells/well) were cultured in 96-well flat-bottom tissue culture plates. After experimental treatments, cells were washed three times with cold phosphate-buffered saline. Cell viability was then assessed using the Cell Count Reagent SF colorimetric assay (Nacalai Tesque, Kyoto, Japan). Briefly, 10 μL of Cell Count Reagent SF was added to each well, followed by incubation at 37°C for 2 hours. Cell viability was quantified by measuring absorbance at 450 nm using a microplate reader (Model 550; Bio-Rad Laboratories, Hercules, CA, USA). The percentage of viable cells was calculated as (T/C) × 100, where T and C represent the mean OD_450_ values of the treated and control groups, respectively.

### 2.5. Cell treatment

HEMn-MPs and G-361 cells were seeded and allowed to attach, after which they were treated with DMSO or staurosporine at the indicated concentrations for the specified durations. Cells were subsequently harvested for protein extraction.

### 2.6. Time-lapse live imaging and cell tracking analysis

HEMn-MPs were cultured in a live-cell imaging chamber (ibiTreat; IBIDI, Martinsried, Germany) and maintained at 37°C with 5% CO_2_. Time-lapse images were acquired every 15 minutes for a total duration of 24 hours using a Biozero 8100 confocal microscope (Keyence Corporation, Osaka, Japan). Melanocytes were identified based on their characteristic dendritic morphology. For cell tracking, time-lapse images were imported into ImageJ software (National Institutes of Health, Bethesda, MD, USA), and individual cell movements were analyzed using the Manual Tracking plugin. Briefly, a melanocyte was selected in the first frame, and its position was recorded in successive frames until the cell exited the field of view or the sequence ended. Tracking data, including cell displacement and trajectories, were exported to Microsoft Excel (Microsoft Corporation, Redmond, WA, USA) for further analysis. Migration distance and velocity were calculated from these data, and cell trajectories were visualized to characterize migration patterns.

### 2.7. Phospho-kinase array

Phosphorylated kinases were profiled using the Proteome Profiler Human Phospho-Kinase Array Kit (R&D Systems, Minneapolis, MN, USA). Procedures were performed following the manufacturer’s protocol with 300 μg of protein lysate per array.

### 2.8. Immunoblot analysis

For protein preparation, cell pellets were extracted as previously described (Yang et al., 2018), and 5 μg of protein were utilized for immunoblot analysis. Primary antibodies were applied at the following dilutions: anti-PARP (CST) 1:1000; anti-cleaved PARP (CST) 1:1000; anti-caspase-3 (CST) 1:1000; anti-cleaved caspase-3 (CST) 1:1000; anti-MITF (Santa Cruz Biotechnology, Dallas, TX, USA) 1:500; anti-TYRP1 (Santa Cruz Biotechnology) 1:500; anti-Melan-A (Abcam, Cambridge, UK) 1:500; anti-PMEL17 (Santa Cruz Biotechnology) 1:500; anti-EDNRB (Abcam) 1:1000; anti-phosphorylated PKCμ (p-PKCμ, Ser744/748; CST) 1:1000; anti-p-PKCμ (Ser916; CST) 1:1000; anti-PKCμ (CST) 1:1000; anti-phosphorylated GSK3α/β (p-GSK3α/β, Ser21/9; CST) 1:1000; anti-GSK3α/β (CST) 1:1000; anti-phosphorylated β-catenin (p-β-catenin, Ser675; CST) 1:1000; anti-β-catenin (CST) 1:1000; anti-CDC42 (CST) 1:500; anti-Rac1/2/3 (CST) 1:500; anti-RhoA (CST) 1:500; anti-RhoB (CST) 1:1000; anti-RhoC (CST) 1:1000; and anti–glyceraldehyde-3-phosphate dehydrogenase (GAPDH; CST) 1:1000. GAPDH served as a loading control.

### 2.9. Immunofluorescence staining of cells

HEMn-MPs and G-361 cells were seeded in six-well plates and treated with DMSO or staurosporine at the indicated concentrations and time points. After treatment, cells were washed thoroughly with ice-cold phosphate-buffered saline and fixed with 4% paraformaldehyde for 5 minutes. Fixed cells were stained with anti-cleaved caspase-3 (CST) or anti-active β-catenin (CST) at a 1:100 dilution. Nuclei were counterstained with Hoechst 33342 (Invitrogen) at a 1:500 dilution. Images were acquired using a Biozero 8100 confocal microscope (Keyence Corporation).

### 2.10. Animals and in vivo experimental procedures

Six male Kwl:JY-4 black guinea pigs (6 weeks of age) were obtained from Tokyo Laboratory Animals Science (Tokyo, Japan) and maintained under specific pathogen-free conditions. After dorsal hair removal with an electric shaver, three or four discrete 1 cm × 1 cm skin areas were designated on each animal and assigned to different experimental conditions. Each experimental group included three guinea pigs. In the normally pigmented skin model, designated areas received daily topical application of vehicle or staurosporine at concentrations of 250, 500, or 1000 μM for 14 consecutive days. In the RD-treated skin model, leukoderma was induced by daily topical application of 30% (*w/v*) RD for 30 days. After cessation of RD treatment, depigmented areas were treated daily with vehicle or staurosporine (1000 μM) for an additional 14 days. Skin samples were collected 24 hours after the final topical treatment. All efforts were made to minimize animal suffering. All procedures were conducted in accordance with the Guiding Principles for the Care and Use of Laboratory Animals and were approved by the Committee for Animal Experiments at Osaka Metropolitan University (Permit No. 18042).

### 2.11. Measurement of skin color

Skin color in treated areas was measured using a portable reflectance spectrophotometer (CM-26d; Konica Minolta, Tokyo, Japan). Color values were expressed as L* values, representing skin lightness on a scale from 0 (black) to 100 (white).

### 2.12. Fluorescent immunohistochemical staining of guinea pig skin tissues

Dorsal skin tissues were fixed in 10% formaldehyde and embedded in paraffin. Paraffin sections (5 μm) were prepared for fluorescent immunohistochemical staining. Sections were deparaffinized, rehydrated, and incubated overnight at 4°C with primary antibodies against TYRP1 (1:200; Sigma-Aldrich) and β-catenin (1:200; CST). After they had been washed, sections were incubated with species-appropriate Alexa Fluor-conjugated secondary antibodies (anti-rabbit IgG Alexa Fluor 488 and anti-mouse IgG Alexa Fluor 555; Invitrogen). Nuclei were counterstained with Hoechst 33342 (1:500; Invitrogen). Images were acquired using a fluorescence microscope or a Biozero 8100 confocal microscope (Keyence Corporation).

### 2.13. Hematoxylin and eosin staining of guinea pig skin tissues

Dorsal skin tissues were fixed in 10% formaldehyde, embedded in paraffin, and sectioned at 5 μm. Sections were deparaffinized in xylene, rehydrated through graded ethanol solutions, and stained with hematoxylin and eosin according to standard protocols. After they had been stained, sections were rinsed with water, dehydrated through graded ethanol, and mounted for imaging.

### 2.14. Statistical analyses

All experiments were performed at least three times. Data are presented as mean ± standard deviation (SD). Statistical analyses were conducted using one-way analysis of variance (ANOVA), followed by a Bonferroni post hoc test, in Microsoft Excel (Microsoft Corporation). A p value < 0.05 was considered statistically significant.

## 3. Results

### 3.1. Staurosporine induces apoptosis in melanoma cells but not normal human melanocytes

To determine whether staurosporine exerts distinct effects in normal melanocytes and melanoma cells, we compared its impact on cell viability and apoptosis in primary normal human epidermal melanocytes (HEMn-MP) and the human melanoma cell line G-361. Cell viability assays revealed a pronounced cell type-specific response to staurosporine treatment (Figure 1A). In G-361 melanoma cells, staurosporine caused a pronounced, dose-dependent reduction in viability, whereas HEMn-MP melanocytes maintained high viability across the same concentration range. Notably, melanocytes exhibited minimal viability loss even at concentrations that were strongly cytotoxic to melanoma cells, indicating robust resistance to staurosporine-induced cell death. Consistent with these findings, phase-contrast microscopy showed profound morphological changes in melanoma cells after staurosporine exposure, including cell shrinkage and loss of adherence; melanocytes displayed no overt signs of cytotoxicity or detachment (Figure 1B). These observations suggested that staurosporine selectively triggers cytotoxic responses in melanoma cells but not normal melanocytes. We next examined whether the differential effects of staurosporine on cell viability were associated with apoptosis. Immunoblot analysis demonstrated robust PARP cleavage and caspase-3 activation in G-361 cells after staurosporine treatment, whereas no detectable cleavage of PARP or caspase-3 was observed in HEMn-MP melanocytes under identical conditions (Figure 1C). These results indicate that staurosporine induces classical caspase-dependent apoptosis in melanoma cells but fails to activate apoptotic signaling in normal melanocytes. To confirm these findings at the single-cell level, we performed immunofluorescence staining for cleaved caspase-3. Staurosporine-treated G-361 cells showed a substantial increase in cleaved caspase-3-positive cells, whereas HEMn-MP melanocytes largely maintained cleaved caspase-3 negativity (Figure 1D). Collectively, these data demonstrate that staurosporine elicits fundamentally distinct cellular responses in malignant versus normal pigment cells, inducing apoptosis in melanoma cells without activating apoptotic signaling in normal melanocytes.

**Figure 1.**
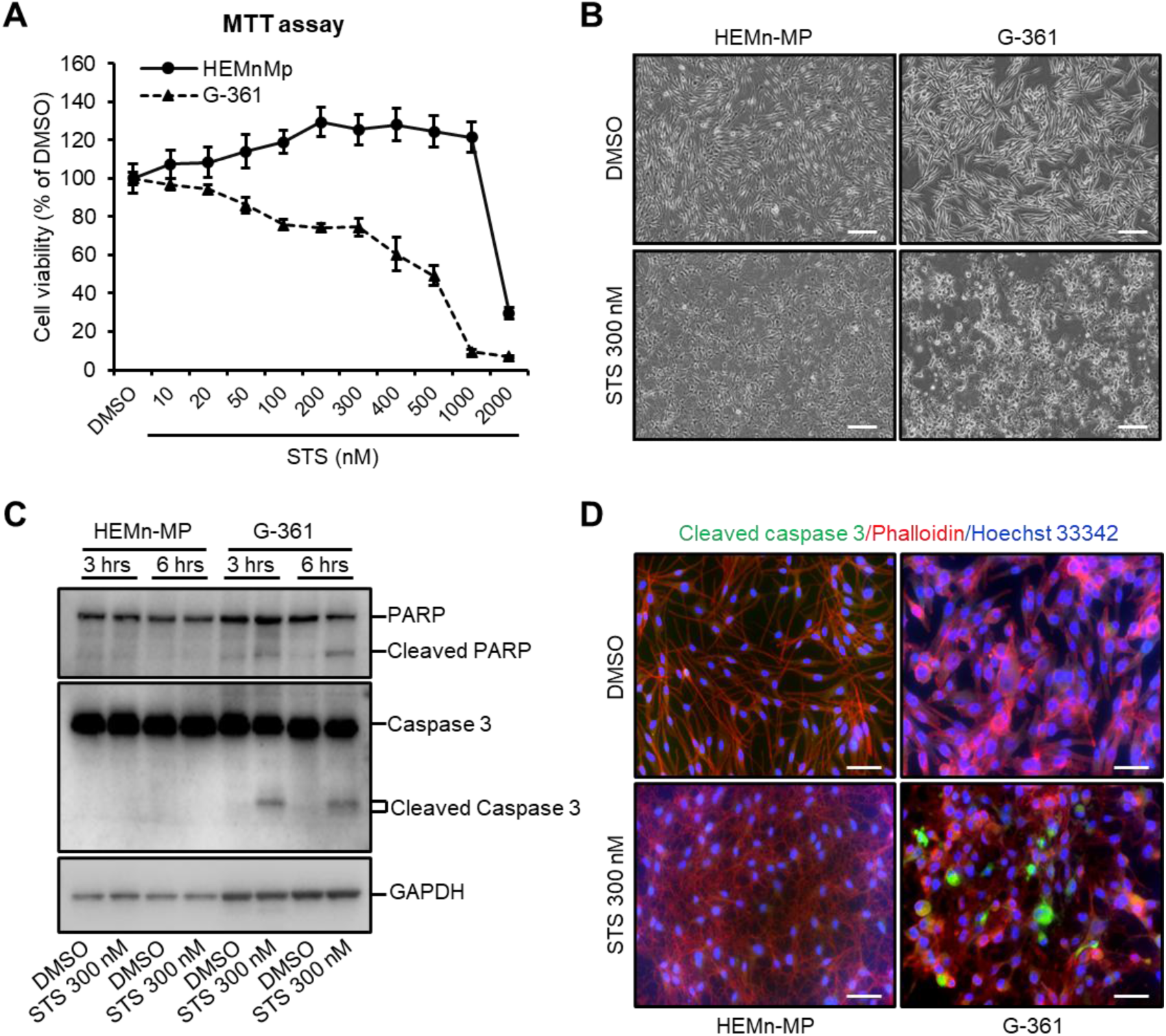
Effects of staurosporine on cell viability and apoptosis in melanocytes and melanoma cells. **(A)** Cell viability of normal human epidermal melanocytes (HEMn-MP) and melanoma cells (G-361) after treatment with increasing concentrations of staurosporine (STS) for 24 hours. Data are presented as percentages of DMSO-treated controls. **(B)** Representative bright-field images of HEMn-MP and G-361 cells treated with DMSO or STS (300 nM), illustrating differences in cell attachment and morphology. Scale bars, 200 μm. **(C)** Immunoblot analysis of PARP, cleaved PARP, caspase-3, and cleaved caspase-3 in HEMn-MP and G-361 cells treated with DMSO or STS for the indicated durations. GAPDH served as a loading control. **(D)** Immunofluorescence staining for cleaved caspase-3 (green) with phalloidin (red) and Hoechst 33342 (blue) in HEMn-MP and G-361 cells treated with DMSO or STS (300 nM). Scale bars, 50 μm.

### 3.2. Staurosporine induces dendrite extension and melanocyte differentiation-associated phenotypes

After establishing that staurosporine does not induce apoptosis in normal melanocytes, we observed pronounced changes in melanocyte morphology. Bright-field imaging and nuclear staining showed that staurosporine-treated melanocytes remained viable and adherent but underwent pronounced morphological remodeling compared with dimethyl sulfoxide (DMSO)-treated controls (Figure 2A; Supplementary Videos 1 and 2). Specifically, staurosporine-treated melanocytes displayed elongated cell bodies and increased dendritic extensions, consistent with enhanced dendricity rather than cytotoxic stress. To characterize the dynamic behavior underlying these changes, we performed time-lapse live-cell imaging and single-cell trajectory analysis. Under control conditions, melanocytes exhibited active migration with extended trajectories, whereas staurosporine-treated melanocytes showed more confined movement with shorter trajectories (Figure 2B, Supplementary Video 3 and 4). Quantitative analysis confirmed a significant increase in dendrite number per cell after staurosporine treatment (Figure 2C), accompanied by a substantial reduction in average migration velocity (Figure 2D). These findings indicate that staurosporine promotes a shift from a motile phenotype to a more stationary, dendrite-rich state characteristic of differentiated melanocytes.

**Figure 2.**
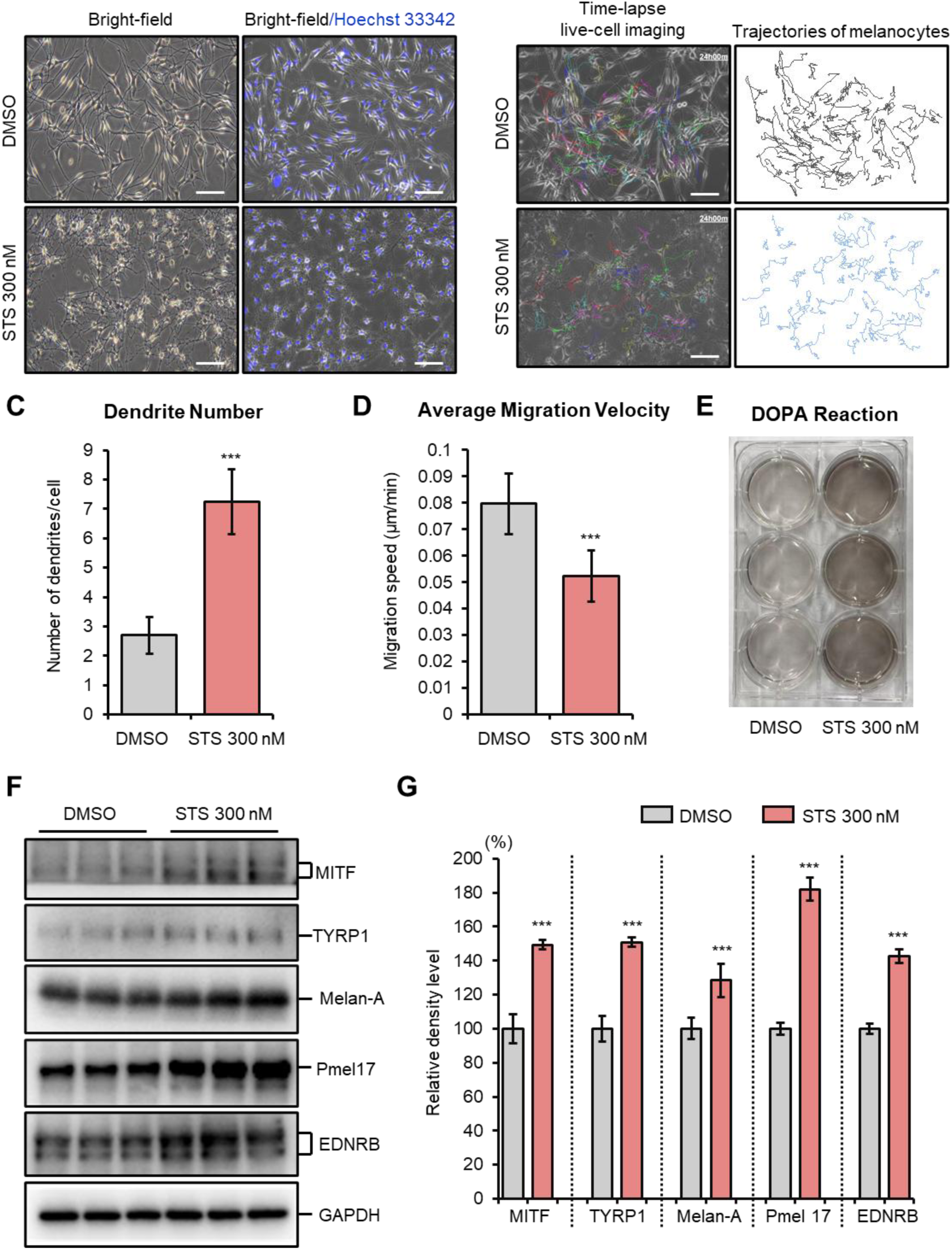
Effects of staurosporine on melanocyte morphology, dendrite formation, and melanogenesis. **(A)** Representative bright-field images of normal human epidermal melanocytes treated with DMSO or staurosporine (STS; 300 nM) for 24 hours. Scale bars, 100 μm. **(B)** Time-lapse live-cell imaging of melanocytes treated with DMSO or STS, acquired at 15-minute intervals over 24 hours. Single-cell tracking analysis was used to assess melanocyte motility. Representative cell trajectories are shown in right panel; dots indicate starting positions. White scale bar, 50 μm. **(C)** Quantification of dendrite number per melanocyte after treatment with DMSO or STS. **(D)** Average migration velocity of melanocytes calculated from single-cell tracking analysis. **(E)** Representative images of DOPA staining in melanocytes treated with DMSO or STS. **(F)** Immunoblot analysis of melanocyte differentiation- and melanogenesis-associated proteins (MITF, TYRP1, Melan-A, PMEL17, and EDNRB) in melanocytes treated with DMSO or STS for 48 hours. GAPDH served as a loading control. **(G)** Quantification of relative protein band intensities shown in **(F)**. Data in **(C)**, **(D)**, and **(G)** represent three independent experiments and are presented as mean ± SD. *** p < 0.01; statistical significance was assessed by one-way ANOVA with a Bonferroni post hoc test.

We next assessed whether these staurosporine-induced morphological and behavioral changes were associated with enhanced melanogenic activity. 3,4-Dihydroxyphenylalanine (DOPA) reaction assays demonstrated increased melanin synthesis in staurosporine-treated melanocytes relative to controls (Figure 2E), indicating functional activation of melanogenesis. Consistent with this observation, immunoblot analysis showed upregulation of key melanocyte differentiation and melanogenic markers, including microphthalmia-associated transcription factor (MITF), tyrosinase-related protein 1 (TYRP1), melanocyte antigen (Melan-A), premelanosome protein (PMEL17), and endothelin receptor type B (EDNRB), after staurosporine treatment (Figure 2F). Densitometric analysis confirmed significant increases in expression of these proteins compared with DMSO-treated cells (Figure 2G). Collectively, these results demonstrate that staurosporine induces coordinated melanocyte functional remodeling, characterized by increased dendricity, reduced migratory behavior, and enhanced melanogenic differentiation. These effects occur in the absence of apoptosis, supporting a model in which staurosporine promotes melanocyte maturation and functional activation rather than cytotoxic stress.

### 3.3 Staurosporine suppresses PKCμ signaling and stabilizes β-catenin in melanocytes

To clarify signaling mechanisms underlying the staurosporine-induced melanocyte phenotypic shift, we first performed an unbiased phospho-kinase profiling analysis. Human phospho-kinase arrays revealed considerable alterations in kinase phosphorylation after staurosporine treatment, including a pronounced reduction in glycogen synthase kinase 3α/β (GSK3α/β) phosphorylation at Ser21/9 compared with DMSO-treated controls (Figure 3A), suggesting modulation of β-catenin regulatory pathways. Staurosporine has long been characterized as a potent, broad-spectrum PKC inhibitor (Yamamoto et al., 1992); however, PKC isoforms targeted by staurosporine in normal human melanocytes have not been defined. Therefore, we systematically examined the phosphorylation status of major PKC isoforms expressed in melanocytes. Immunoblot analysis showed that staurosporine treatment significantly reduced PKCμ phosphorylation at Ser744/748 and Ser916, consistent with suppression of PKC signaling activity (Figure 3B, 3C). In contrast, phosphorylation of other PKC isoforms, including PKCα, PKCβII, PKCζ/λ, PKCθ, and PKCδ, remained largely unchanged under the same conditions (Supplementary Figure 1), indicating that staurosporine has a preferential effect on PKCμ signaling in melanocytes. In parallel, GSK3α/β phosphorylation at Ser21/9 was greatly decreased, suggesting altered regulation of the GSK3-dependent β-catenin destruction complex. Concomitantly, β-catenin phosphorylation at Ser675 was significantly reduced (Figure 3B, 3C). To determine whether suppression of PKC signaling by staurosporine led to accumulation of transcriptionally active β-catenin, we performed immunofluorescence staining for active β-catenin. Whereas control melanocytes showed weak and diffuse staining, staurosporine-treated melanocytes exhibited a pronounced increase in active β-catenin-positive cells, with prominent nuclear localization (Figure 3D). Our findings indicate β-catenin-dependent signaling activation downstream of PKC inhibition. Taken together, these data demonstrate that staurosporine suppresses PKC signaling in melanocytes, leading to GSK3α/β activity inhibition and active β-catenin stabilization. This reprogrammed signaling provides a mechanistic basis for the staurosporine-induced changes in melanocyte morphology and differentiation shown in Figure 2.

**Figure 3.**
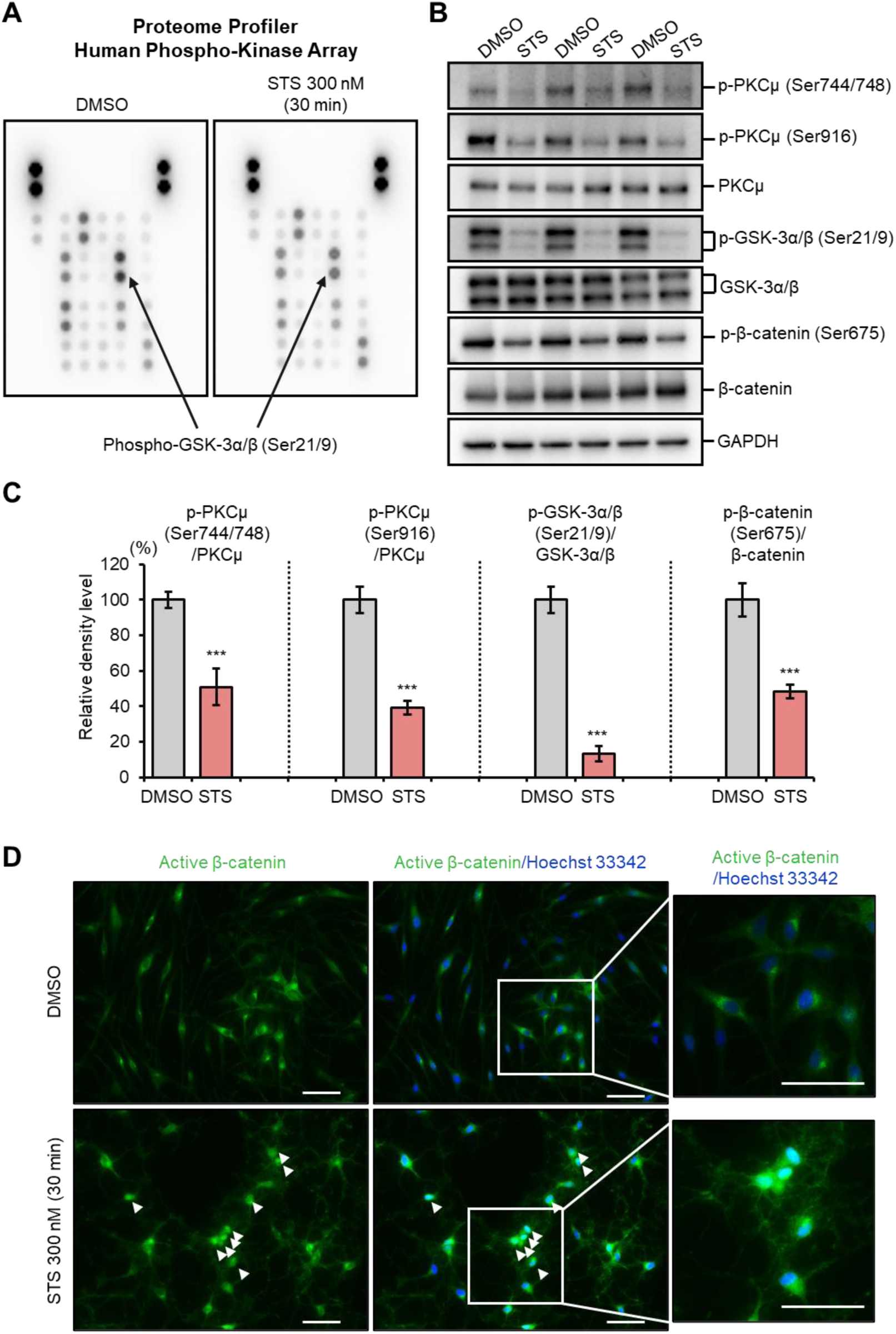
Phospho-kinase profiling and analysis of staurosporine-induced signaling pathways in melanocytes. **(A)** Human phospho-kinase array analysis of melanocytes treated with DMSO or staurosporine (STS; 300 nM) for 30 minutes. Representative array images and quantification of selected phosphorylated kinases are shown. **(B)** Immunoblot analysis of PKD/PKCμ phosphorylation (Ser744/748 and Ser916), total PKD/PKCμ, GSK3α/β phosphorylation (Ser21/9), total GSK3α/β, β-catenin phosphorylation (Ser675), and total β-catenin in melanocytes treated with DMSO or STS for 30 minutes. **(C)** Quantification of relative protein band intensities shown in **(B)**, normalized to corresponding total protein levels or GAPDH, as indicated. **(D)** Immunofluorescence staining for active β-catenin in melanocytes treated with DMSO or STS. Nuclei were counterstained with Hoechst 33342. Representative active β-catenin-positive cells are indicated by white arrowheads and shown at higher magnification in right panel. Scale bars, 100 μm. Data in **(C)** represent three independent experiments and are presented as mean ± SD. *** p < 0.01; statistical significance was assessed by one-way ANOVA with a Bonferroni post hoc test.

### 3.4. Differential regulation of Rho GTPase expression underlies staurosporine-induced melanocyte dendricity

To investigate the molecular basis of staurosporine-induced morphological changes, we focused on the Rho family of small GTPases, which are central regulators of actin cytoskeleton dynamics and dendrite formation in melanocytes (G. Scott, 2002). Immunoblot analysis demonstrated that staurosporine treatment (300 nM) significantly increased protein levels of cell division cycle 42 (CDC42) and Ras-related C3 botulinum toxin substrate 1/2/3 (Rac1/2/3), while concomitantly reducing RhoA expression (Figure 4A, 4C). In contrast, expression patterns of Ras homolog family member B (RhoB) and Ras homolog family member C (RhoC) remained largely unchanged, indicating selective modulation of specific Rho GTPase family members.

**Figure 4.**
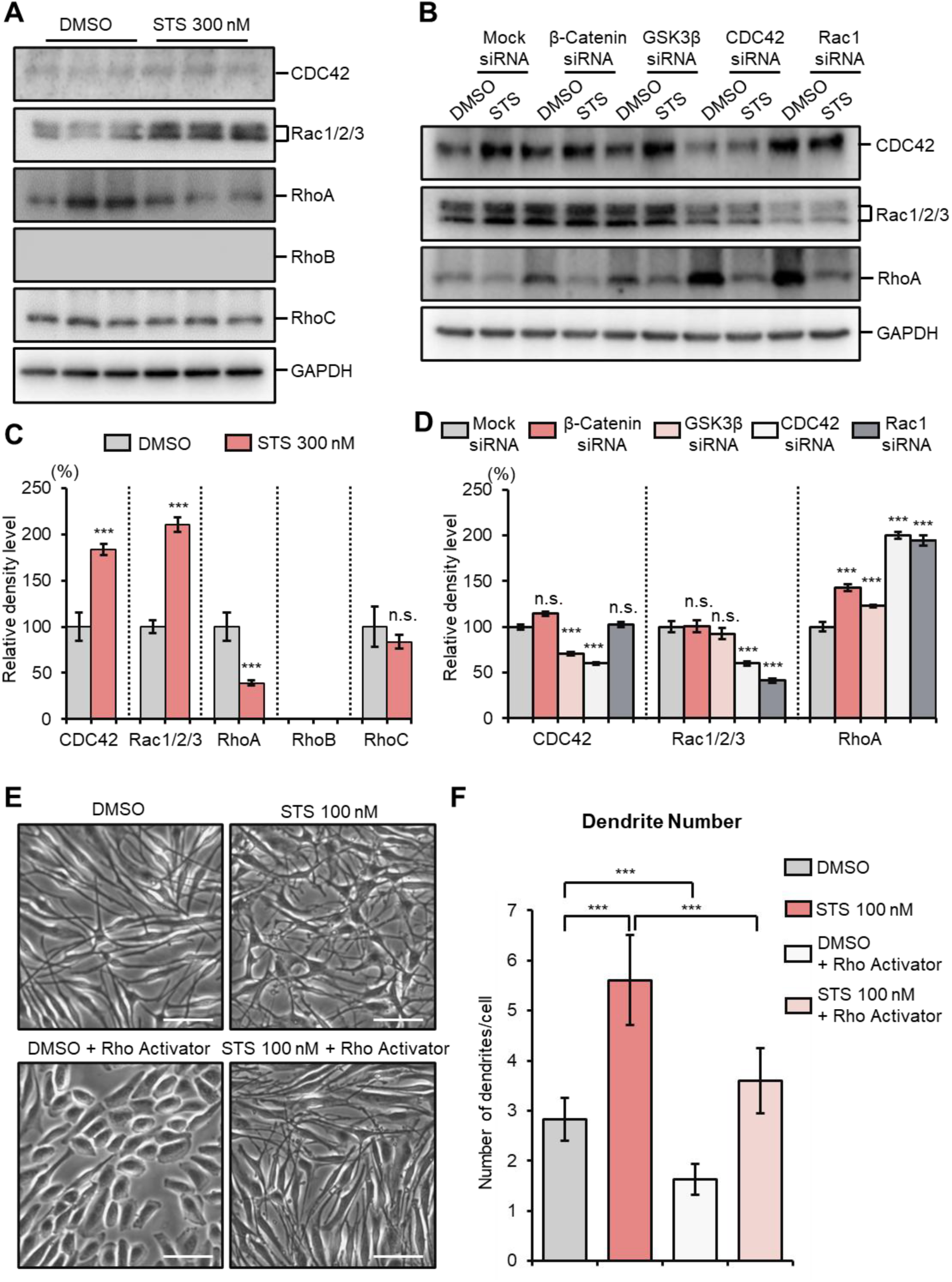
Alterations in RhoA and cytoskeleton-associated protein expression in melanocytes after staurosporine treatment. **(A)** Immunoblot analysis of CDC42, Rac1/2/3, RhoA, RhoB, and RhoC protein levels in normal human epidermal melanocytes treated with DMSO or staurosporine (STS; 300 nM) for 48 hours. GAPDH served as a loading control. **(B)** Immunoblot analysis of CDC42, Rac1/2/3, and RhoA protein levels in melanocytes 24 hours after transfection with mock control siRNA or siRNAs targeting β-catenin, GSK3β, CDC42, or Rac1, followed by treatment with DMSO or STS for 24 hours. GAPDH served as a loading control. **(C)** Densitometric quantification of protein levels shown in (**A**), normalized to GAPDH and expressed relative to the DMSO-treated group. **(D)** Densitometric quantification of protein levels shown in **(B)**, normalized to GAPDH and expressed relative to the mock siRNA group. **(E)** Representative bright-field images of melanocytes treated with DMSO or staurosporine (STS; 100 nM) for 24 hours in the absence or presence of a Rho activator (5 μg/mL). Scale bars, 50 μm. **(F)** Quantification of dendrite number per melanocyte under the conditions shown in **(E)**. Data in **(C)**, **(D)**, and **(F)** represent three independent experiments and are presented as mean ± SD. *** p < 0.001; n.s., not significant. Statistical significance was assessed by one-way ANOVA with a Bonferroni post hoc test.

To define upstream regulatory relationships governing RhoA expression, we transfected melanocytes with small interfering RNAs (siRNAs) targeting β-catenin, GSK-3β, CDC42, or Rac1. Knockdown of β-catenin or GSK-3β resulted in a pronounced increase in RhoA protein levels compared with control siRNA, indicating that GSK-3β and β-catenin signaling function upstream to restrain RhoA expression (Figure 4B, 4D). Notably, depletion of CDC42 or Rac1 led to pronounced upregulation of RhoA, supporting a model in which both CDC42 and Rac1 act as upstream suppressors of RhoA expression in melanocytes. Consistent with this regulatory hierarchy, pharmacologic activation of RhoA antagonized dendritic extension and reduced dendrite number per cell (Figure 4E, 4F), supporting a central role for RhoA suppression in dendrite formation. Collectively, these findings indicate that coordinated suppression of RhoA expression by upstream β-catenin, CDC42, and Rac1 is a key molecular feature associated with enhanced melanocyte dendricity.

### 3.5. Topical staurosporine enhances epidermal pigmentation and increases melanocyte abundance in guinea pig skin

To determine whether the pro-differentiation and dendrite-promoting effects observed *in vitro* translate to an *in vivo* setting, we assessed the effects of topical staurosporine on guinea pig skin pigmentation. Dorsal skin areas were shaved and treated daily with vehicle or increasing concentrations of staurosporine for 14 days (Figure 5A). Macroscopic examination revealed dose-dependent darkening of treated skin after topical staurosporine application, as evidenced by digital photography, wood lamp examination, and dermoscopic analysis (Figure 5B). Quantitative colorimetric assessment confirmed a significant, dose-dependent decrease in skin color L* values in staurosporine-treated sites compared with vehicle controls, indicating enhanced pigmentation (Figure 5C). Histological and immunofluorescence analyses further supported these findings. TYRP1 immunostaining demonstrated a substantial increase in the number of TYRP1-positive melanocytes along the basal epidermis in staurosporine-treated skin compared with vehicle-treated controls (Figure 5D, 5E). Hematoxylin and eosin staining showed preserved epidermal architecture without overt tissue damage or inflammation, indicating good local tolerability of topical staurosporine treatment (Figure 5D). Consistent with the *in vitro* findings, staurosporine-treated skin exhibited a pronounced increase in nuclear β-catenin localization within epidermal melanocytes, accompanied by enhanced TYRP1 expression (Figure 5F). These results indicate that topical staurosporine promotes melanocyte differentiation and increases melanocyte abundance within the epidermis in association with β-catenin-linked signaling pathways. Collectively, our observations suggest that topical staurosporine enhances epidermal pigmentation and increases melanocyte abundance in guinea pig skin, supporting its translational potential as a topical modulator of melanocyte function.

**Figure 5.**
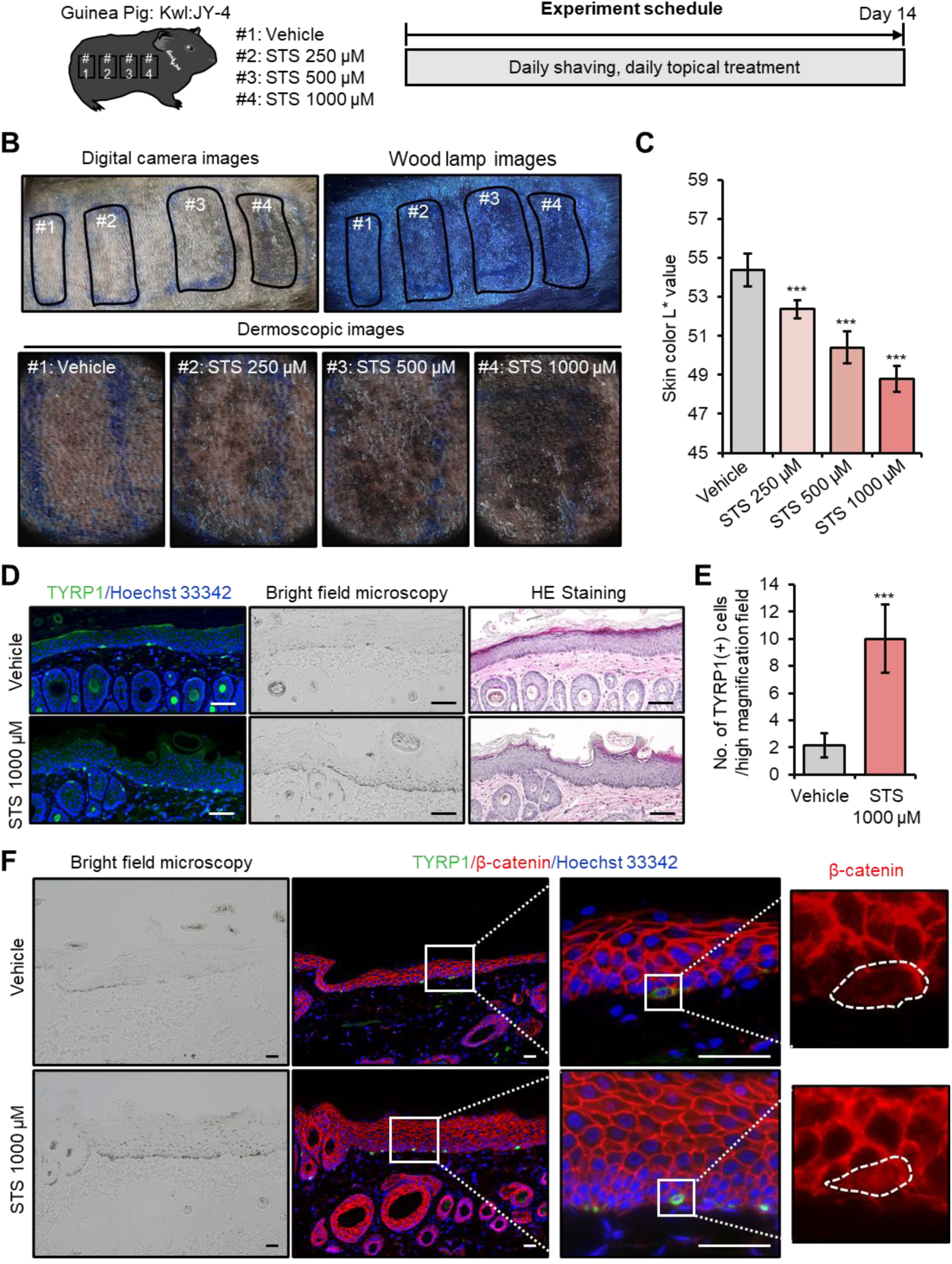
Topical staurosporine promotes skin pigmentation and epidermal melanocyte accumulation in guinea pigs. **(A)** Experimental schedule. **(B)** Representative digital photographs, Wood lamp images, and dermoscopic images of treated skin areas after 14 days of topical application. **(C)** Quantification of skin color L* values for the treated areas shown in **(B)**. **(D)** Representative images of TYRP1 immunofluorescence (green) with Hoechst 33342 nuclear counterstaining (blue), bright-field microscopy, and hematoxylin and eosin staining of skin sections from vehicle- and STS-treated areas (1000 μM). White scale bar, 100 μm. **(E)** Quantification of TYRP1-positive cells per high-power field. **(F)** Representative bright-field images and immunofluorescence staining for TYRP1 (green), β-catenin (red), and nuclei (Hoechst 33342; blue) in skin sections from vehicle- and STS-treated sites (1000 μM). Enlarged views highlight β-catenin localization in epidermal melanocytes (outlined by dashed lines). White scale bar, 50 μm. Data in **(C)** and **(E)** are presented as mean ± SD. *** p < 0.01; statistical significance was assessed by one- way ANOVA with a Bonferroni post hoc test.

### 3.6. Topical staurosporine promotes repigmentation and epidermal melanocyte recovery in RD-treated guinea pig skin

To assess the effects of topical staurosporine in depigmentation disorders, we used an RD-induced leukoderma guinea pig model, which recapitulates chemical-induced melanocyte loss and depigmentation *in vivo* (Inoue et al., 2021; Kuroda, Takahashi, Sakaguchi, Matsunaga, & Suzuki, 2014). Repeated topical application of 30% RD for 30 days resulted in obvious skin depigmentation (Figure 6A, 6B). After cessation of RD treatment, slight spontaneous recovery of skin pigmentation and epidermal melanocytes was observed over the subsequent 14 days. In contrast, topical application of staurosporine (1000 μM) markedly enhanced repigmentation in RD-treated skin and significantly attenuated RD-induced skin lightening compared with vehicle-treated controls (Figure 6B, 6C). Immunohistochemical and immunofluorescence analyses also demonstrated increased numbers of epidermal melanocytes in staurosporine-treated areas relative to vehicle-treated RD skin (Figure 6D, 6E). Finally, epidermal melanocytes in staurosporine-treated skin displayed increased nuclear localization of β-catenin compared with RD-treated controls (Figure 6F). These findings indicate that topical staurosporine promotes repigmentation accompanied by epidermal melanocyte recovery in RD-treated guinea pig skin.

**Figure 6.**
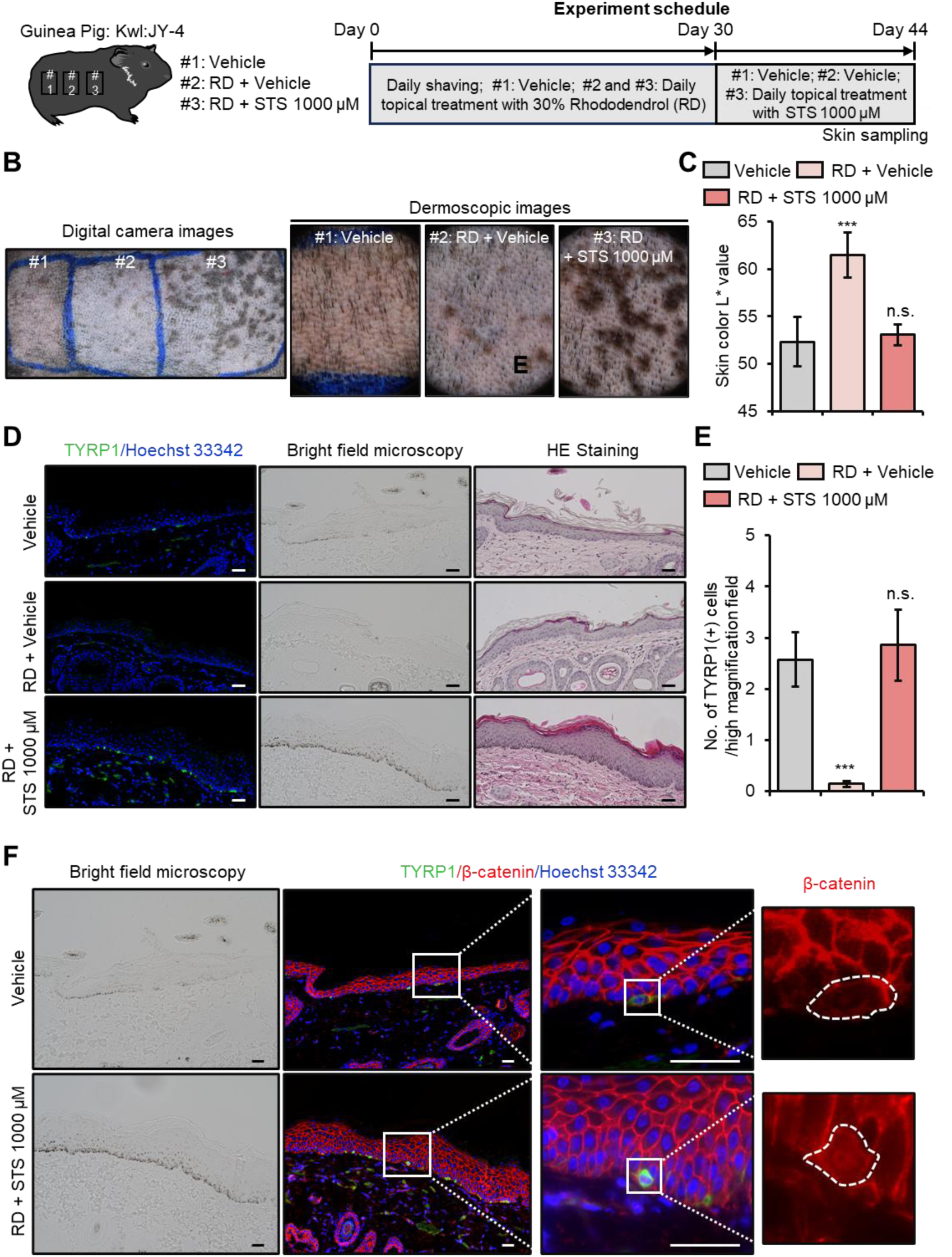
Topical staurosporine enhances repigmentation in guinea pigs with rhododendrol-induced depigmentation. (**A**) Experimental schedule. (**B**) Representative digital photographs and dermoscopic images of skin areas under the indicated treatments. (**C**) Quantification of skin color L* values in the indicated treatment groups. (**D**) Representative immunofluorescence staining for TYRP1 (green) with Hoechst 33342 nuclear counterstaining (blue), together with bright-field microscopy and hematoxylin and eosin staining of skin sections from each group. White scale bar, 50 μm. (**E**) Quantification of TYRP1-positive cells per high-power field. **(F)** Representative bright-field and immunofluorescence images showing TYRP1 (green), β-catenin (red), and nuclear counterstaining (Hoechst 33342; blue). Enlarged views highlight epidermal melanocytes and β-catenin localization. Data in **(C)** and **(E)** are presented as mean ± SD. *** p < 0.001; n.s., not significant. Statistical significance was assessed by one-way ANOVA with a Bonferroni post hoc test.

## 4. Discussion

Staurosporine has long been recognized as a potent, broad-spectrum kinase inhibitor and a canonical inducer of apoptosis in melanoma and other malignant cells (Gescher, 1998; Omura et al., 1995; Yamamoto et al., 1992). Beyond its cytotoxic effects, accumulating evidence from non-melanocytic systems indicates that, at nanomolar and sub-cytotoxic concentrations, staurosporine can promote cellular differentiation and profound morphological remodeling (Borge et al., 1997; Kronfeld, Zsukerman, Kazimirsky, & Brodie, 1995; Kulyk, 1991; Rasouly et al., 1996; Sano et al., 1994; Thompson & Levin, 2010). Such effects have been particularly well documented in neuronal and mesenchymal lineages, where staurosporine induces neurite outgrowth, cytoskeletal reorganization, and lineage-specific maturation. Despite these observations, the impact of staurosporine on normal human melanocyte biology has remained largely unexplored.

In this study, we demonstrate that staurosporine elicits fundamentally distinct, cell type-specific responses in normal melanocytes compared with melanoma cells. While melanoma cells undergo apoptosis in response to staurosporine, primary human melanocytes display marked resistance to cell death and instead activate a robust maturation program. At sub-cytotoxic concentrations, staurosporine promotes dendritic elongation, enhances melanogenic activity, and increases functional pigmentation in vitro and in vivo. These findings highlight the intrinsic capacity of melanocytes to interpret kinase inhibition signals in a differentiation-promoting rather than cytotoxic manner.

Our data suggest that inhibition of PKC-dependent signaling relieves constraints on melanocyte maturation, enabling coordinated regulation of pigmentation and cell morphology. This response is associated with stabilization and nuclear accumulation of β-catenin, a key regulator of melanocyte differentiation and lineage maintenance (Gallagher et al., 2013; Larue, Kumasaka, & Goding, 2003; Yamada et al., 2013), as well as with altered activity of Rho family GTPases that govern actin cytoskeletal dynamics (Lacour, Gordon, Eller, Bhawan, & Gilchrest, 1992; G. Scott, 2002; G. A. Scott & Cassidy, 1998). Although β-catenin is classically activated through canonical Wnt signaling (Yamada et al., 2013), our findings indicate that, in melanocytes, β-catenin-dependent differentiation can be engaged through a non-canonical, Wnt-independent mechanism. This distinction underscores the context-dependent roles of β-catenin signaling in pigment cells, where it favors terminal differentiation and reduced motility rather than proliferation.

Importantly, the coordinated activation of melanogenic gene expression and dendritic remodeling provides a structural and functional framework for efficient pigment production and melanosome transfer. By simultaneously enhancing the biosynthetic capacity of melanocytes and the dendritic network required for pigment delivery, staurosporine engages a comprehensive maturation program that mirrors key features of physiological pigmentation.

The physiological relevance of this response was supported by *in vivo* experiments demonstrating increased pigmentation following topical staurosporine application in guinea pig skin, in the absence of overt inflammation or tissue damage. Moreover, staurosporine accelerated repigmentation in a rhododendrol-induced leukoderma model, characterized by restoration of epidermal melanocytes and increased nuclear β-catenin localization. These observations indicate that activation of melanocyte-intrinsic maturation pathways is sufficient to enhance pigmentation under both normal and depigmented conditions.

The differential sensitivity of normal melanocytes and melanoma cells to staurosporine-induced apoptosis highlights fundamental differences in kinase network regulation between benign and malignant pigment cells. Whereas oncogenic rewiring in melanoma predisposes cells to apoptosis upon kinase inhibition, normal melanocytes appear capable of maintaining cellular homeostasis and redirecting these signals toward differentiation. In contrast to ultraviolet radiation-dependent pigmentation, which relies heavily on keratinocyte-derived paracrine factors (Gordon, Mansur, & Gilchrest, 1989; Halaban et al., 1988; Tomita, Iwamoto, Masuda, & Tagami, 1987; Yaar, Grossman, Eller, & Gilchrest, 1991), the pathway identified here represents an intrinsic, ultraviolet-independent mechanism that directly links cytoskeletal remodeling with melanocyte maturation.

Several limitations should be acknowledged. Although our findings support a predominantly melanocyte-intrinsic mechanism, potential contributions from the epidermal microenvironment, including keratinocyte-derived signals and neurocutaneous interactions (Hara et al., 1996; Kippenberger, Bernd, Bereiter-Hahn, Ramirez-Bosca, & Kaufmann, 1998), cannot be excluded. Future studies will be required to determine whether staurosporine also influences these extrinsic factors to indirectly support melanocyte maturation.

In summary, our study identifies staurosporine as a potent modulator of melanocyte dendricity and pigmentation and reveals a non-canonical maturation program that couples cytoskeletal remodeling to melanogenic activity. These findings provide new insight into the intrinsic regulatory mechanisms governing melanocyte function and highlight dendritic remodeling as a central determinant of effective pigmentation.

## Acknowledgments

The authors thank Kumiko Mitsuyama for administrative support.

