## Supplementary Information for "Staurosporine drives non-canonical melanocyte maturation by coupling β-catenin signaling to actin-dependent dendrite remodeling"

### Supplementary materials & methods

#### *1. Western blot analysis*

A 5-μg aliquot of the extracted proteins was utilized for western blot analysis. Primary antibodies were employed at the following dilutions: anti-p-PKCμ (Ser916) (CST, Danvers, MA, USA) at 1:1000, anti-p-PKCμ (Ser744/748) (CST) at 1:1000, anti-PKCμ (CST) at 1:1000, anti-p-PKCα/βII (Thr638/641) (CST) at 1:1000, anti-PKCα (CST) at 1:1000, anti-p-PKCζ/λ (Thr410/403) (CST) at 1:1000, anti-PKCζ (CST) at 1:1000, anti-p-PKCδ (Thr505) (CST) at 1:1000, anti-p-PKCδ/θ (Ser643/676) (CST) at 1:1000, anti-p-PKCθ (Thr538) (CST) at 1:1000, anti-PKCδ (CST) at 1:1000, and anti-GAPDH (CST) at 1:1000. The anti-GAPDH antibody was employed as a loading control.

### Supplementary figure legends

**Supplementary Figure 1. Effects of staurosporine on PKC isoform phosphorylation in cultured human epidermal melanocytes.** Representative Western blot analysis showing the phosphorylation status and total protein levels of the indicated PKC isoforms in melanocytes treated with 300 nM staurosporine for the indicated time periods. Total protein levels of PKCμ, PKCα, PKCζ, and PKCδ are shown as indicated. GAPDH served as a loading control.

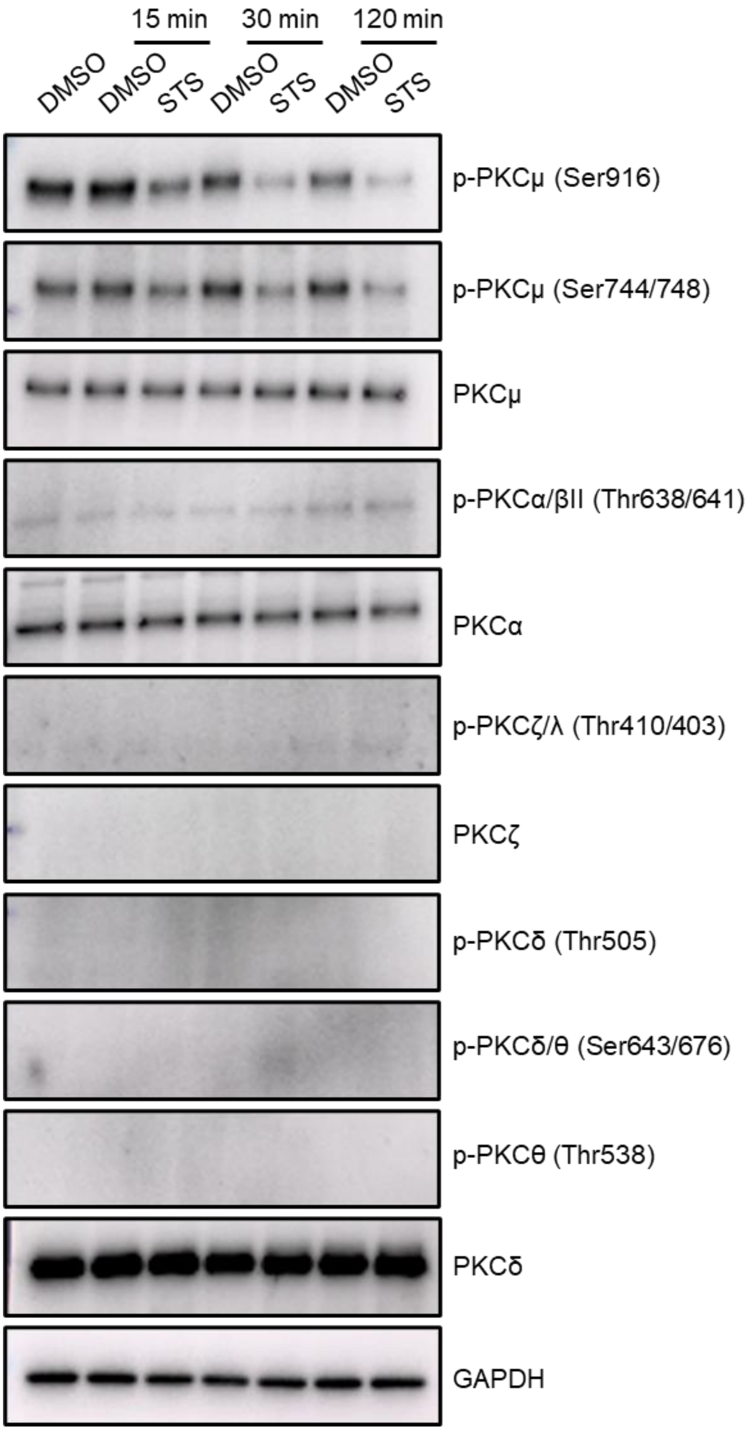


### Supplementary video legends

**Supplementary Video 1:** Time-lapse imaging of cultured HEMn-MPs with dimethyl sulfoxide

(DMSO) treatment as the control.

**Supplementary Video 2:** Time-lapse imaging of cultured HEMn-MPs with 300 nM staurosporine treatment.
